# Peste des Petits Ruminants virus virulence is associated with an early inflammatory profile in the tonsils and cell cycle arrest in lymphoid tissue

**DOI:** 10.1101/2024.04.23.590699

**Authors:** Roger-Junior Eloiflin, Llorenç Grau-Roma, Vincent Lasserre, Sylvie Python, Stephanie Talker, Philippe Totte, Obdulio García- Nicolás, Artur Summerfield, Arnaud Bataille

**Author notes:** Contributed equally (Artur Summerfield, Arnaud Bataille).

## Abstract

Using a systems immunology approach, this study comprehensively explored the immunopathogenesis of Peste des Petits Ruminants (PPR) focussing on strain-dependent differences in virulence. Saanen goats were infected either with the highly virulent Morroco 2008 (MA08) or the low virulent Ivory Coast 1989 (IC89) strain of PPR virus (PPRV). As expected, MA08-infected goats exhibited higher clinical scores, pronounced lymphocyte depletion, and lesions affecting mucosal and lymphoid tissues. CD4 T cells were found to be most affected in terms of depletion and infection in the peripheral blood. Transcriptional analyses of the blood and lymphoid tissue demonstrated activation of interferon type I (IFN-I) responses at three days post infection (dpi) only with MA08, but comparable IFN-I expression levels with MA08 and IC89 at 6 dpi. In contrast, only the MA08 strain induced strong inflammatory and myeloid cell-related transcriptional responses which as observed in tonsils but not in the mesenteric lymph node. This inflammatory response in the tonsil was associated with an extensive damage and infection of the tonsillar epithelium in the crypts, pointing on a barrier defect as a possible cause of inflammation. The other prominent effect induced by MA08, but not IC89, was a strong and early downregulation of cell cycle gene networks in lymphoid tissues. This effect was found in the blood compartment and all analysed lymphoid tissues and can be interpreted as suppressed lymphocyte proliferation that may cause immunosuppression during the first week following MA08 infection. A proteome analysis confirmed elevated synthesis of IFN-I response proteins during infection with both strains, but only the MA08 strain additionally upregulated ribosomal and inflammation-related proteins. In conclusion, the present comprehensive investigation delineates strain-dependent differences in early immunopathological processes associated with severe inflammation disease and a blunted lymphocyte proliferation. Understanding such strain-specific differences is relevant for effective PPRV surveillance strategies.

**Author summary:** Field observations show that the severity of infection with Peste des Petits Ruminants virus (PPRV) is highly dependent on the viral strains and the host infected, but the mechanisms behind these variations are not well understood. Here we compare immune response in Saanen goats infected with high (MA08) and low (IC89) virulent PPRV strains. Analyses revealed a differential immune response: early activation of type I interferon (IFN-I) responses only with MA08, but comparable IFN-I expression levels with MA08 and IC89 at later stages. Additionally, only the MA08 strain triggered inflammatory and myeloid cell-related responses in the tonsils, as well as a disseminated early and marked suppression of lymphocyte proliferation evidenced by cell cycle arrest. CD4 T cells were found to be most affected in terms of depletion in the peripheral blood. Massive infection of the tonsils, particularly for the highly virulent strains, seems to induce epithelial lesions that promotes the inflammatory responses. These findings underscore the importance of understanding strain-specific differences for appropriate surveillance and control of PPR.

## Introduction

In the realm of infectious diseases affecting domestic small ruminants and wild artiodactyls, Peste des Petits Ruminants (PPR) remains an ubiquitous and devastating threat, presenting a significant concern for the global economy, food security and biodiversity[1–3]. The causative pathogen is the PPR virus (PPRV), a member of the Morbillivirus genus, capable of inducing acute and sub-acute clinical manifestations of the disease. PPRV is mainly transmitted through direct contact between healthy and infected animals, or by contaminated aerosols. Indirect transmission may also occur through contaminated feed, water, and fomites, although the importance of such transmission is not yet well understood [4].

The susceptibility of host species to PPRV infection is characterized by marked heterogeneity, with symptoms observed and disease outcome dependent on the virus strain and the host species and breed infected [5]. This heterogeneity is of significant epidemiological relevance and intimately tied to the complex interplay between the host organism and the specific viral strains involved [6,7], but it is unclear how this is associated with differences in innate and adaptive immune responses. Availability of validated diagnostic tests as well as cheap and efficient vaccines are essential to FAO and WOAH efforts to eradicate PPR by 2030[8,9].

Nevertheless, efficiency of surveillance and control strategies are also dependent on improving our understanding of host susceptibility and pathogenesis.

The receptor for PPRV is the Signaling Lymphocyte Activation Molecule (SLAM), as with other morbilliviruses [10], which is mainly expressed by activated lymphocytes. PPRV infection of lymphocytes may prevent clonal expansion of virus-specific lymphocytes thereby suppressing the development of adaptive immune responses, as demonstrated for other morbilliviruses[11–16]. On the other hand, several innate immune mechanisms such as autophagy[17], inflammasome[18], induction of the type-I interferon (IFN-I) response[19], inhibition of nucleotide biosynthesis[20] and apoptosis[21] have been shown *in vitro* to help limit the replication of the virus. Nevertheless, innate mechanisms, particularly virus-induced inflammation, may also play a role in strain-dependent differences in virulence^23^.

To understand these differences in PPRV virulence, we have established a model using Saanen goats. In this model the PPRV strain Morocco 2008 (MA08) induced severe disease, lymphocyte depletion, prominent macroscopic and histological lesions, high viremia, and high virus shedding during the acute phase of disease, whereas the strain Ivory Coast 1989 (IC89) did not [22]. A first *in vitro* experiment had already shown that differential immune response could be observed in peripheral blood mononuclear cells (PMBCs) in this challenge model^23^. Considering the importance of the immune response in morbillivirus pathogenesis, the present study aimed to characterize *in vivo* the differences in the induction of innate immune responses and the initiation of adaptive immune responses in the peripheral blood and lymphoid tissues between MA08 and IC89. To this end, we used blood samples collected during a first experiment^7^ and we performed a second experiment using the same model to collect secondary lymphoid tissues. These are expected to be the primary target of PPRV at early time points post-infection, in order to understand the relationship between innate immune responses and pathogenesis. To achieve this aim, we employed a multi-omics approach including flow cytometry, RNA sequencing and mass spectrometry. Our data demonstrate that virulence is associated with a dysregulated inflammatory response and prominent suppression of cell proliferation in lymphoid tissue.

## Results

### Viral RNA and nucleoprotein in organs and cells

Quantification of viral RNA reads in RNA sequencing data of PBMCs obtained from the first goat experiment (experiment layout depicted in **Fig 1A**) revealed the presence of viral gene expression already at three days post infection (dpi) in animals infected with strain MA08, with clearly increased levels at 7 dpi and decreasing levels at 12 dpi (**Fig 2A**). In contrast, samples from IC89-infected animals had lower viral loads that were only detected at 7 and 12 dpi (**Fig 2A**). Different organs and cells collected during the second animal experiment were investigated in a similar manner for the presence of viral RNA (experiment layout depicted in **Fig 1B**). This confirmed the presence of viral RNA in sampled organs from MA08-infected animals already at 3 dpi with a further increase at 6 dpi (**Fig 2B**). In contrast, in IC89-infected animals viral RNA was detected in only one organ of one animal at 3 dpi (**Fig 2B**). In all compartments, MA08-infected animals had the highest number of viral readings. Our data also indicate a stronger viral presence in the tonsils than in other organs tested for both viral strains (**Fig 2B**; see also confirmatory RT-qPCR for viral RNA, **S1 Fig**).

**Fig 1.**
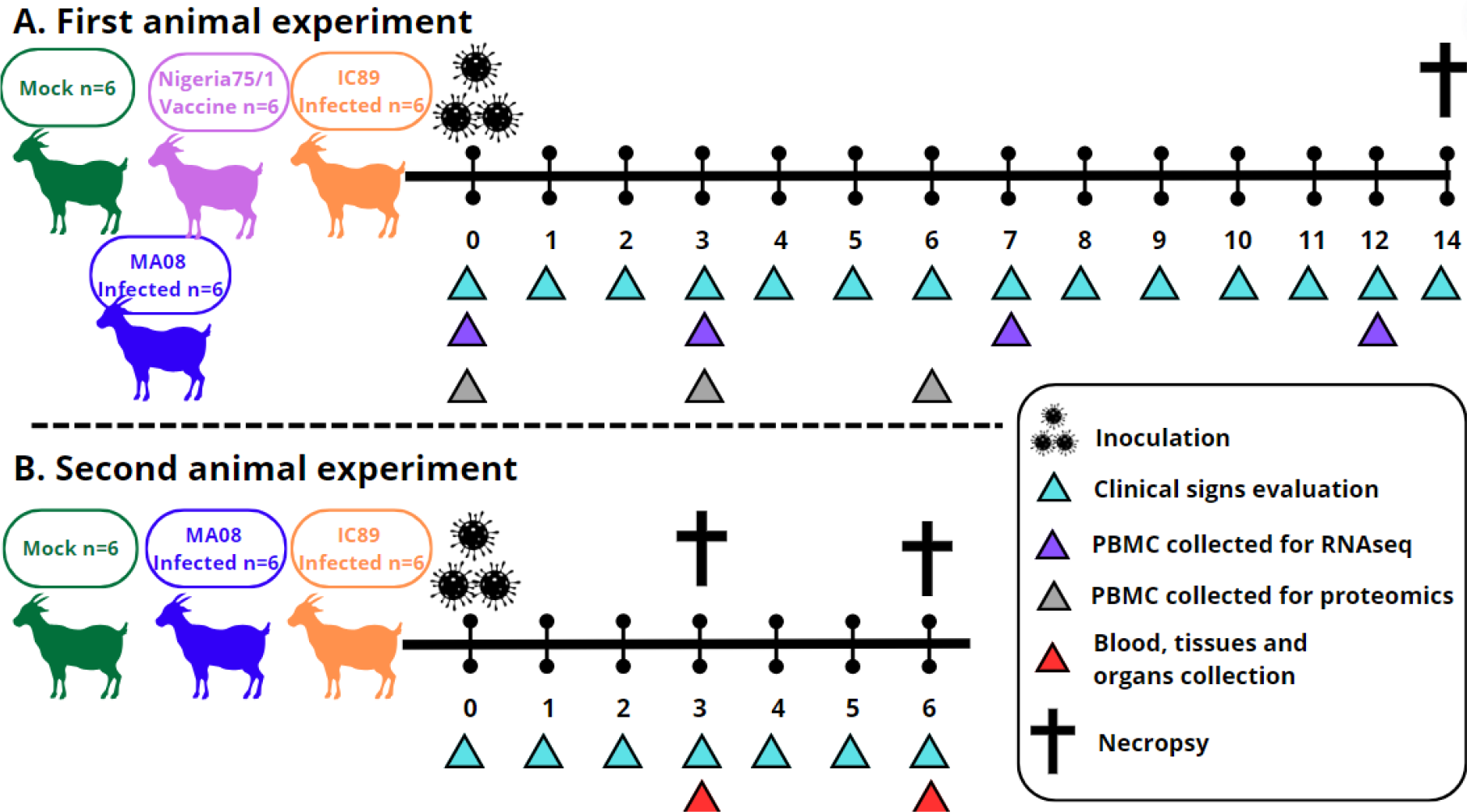
Graphical summary of animal experiments included in this study.

**Fig 2.**
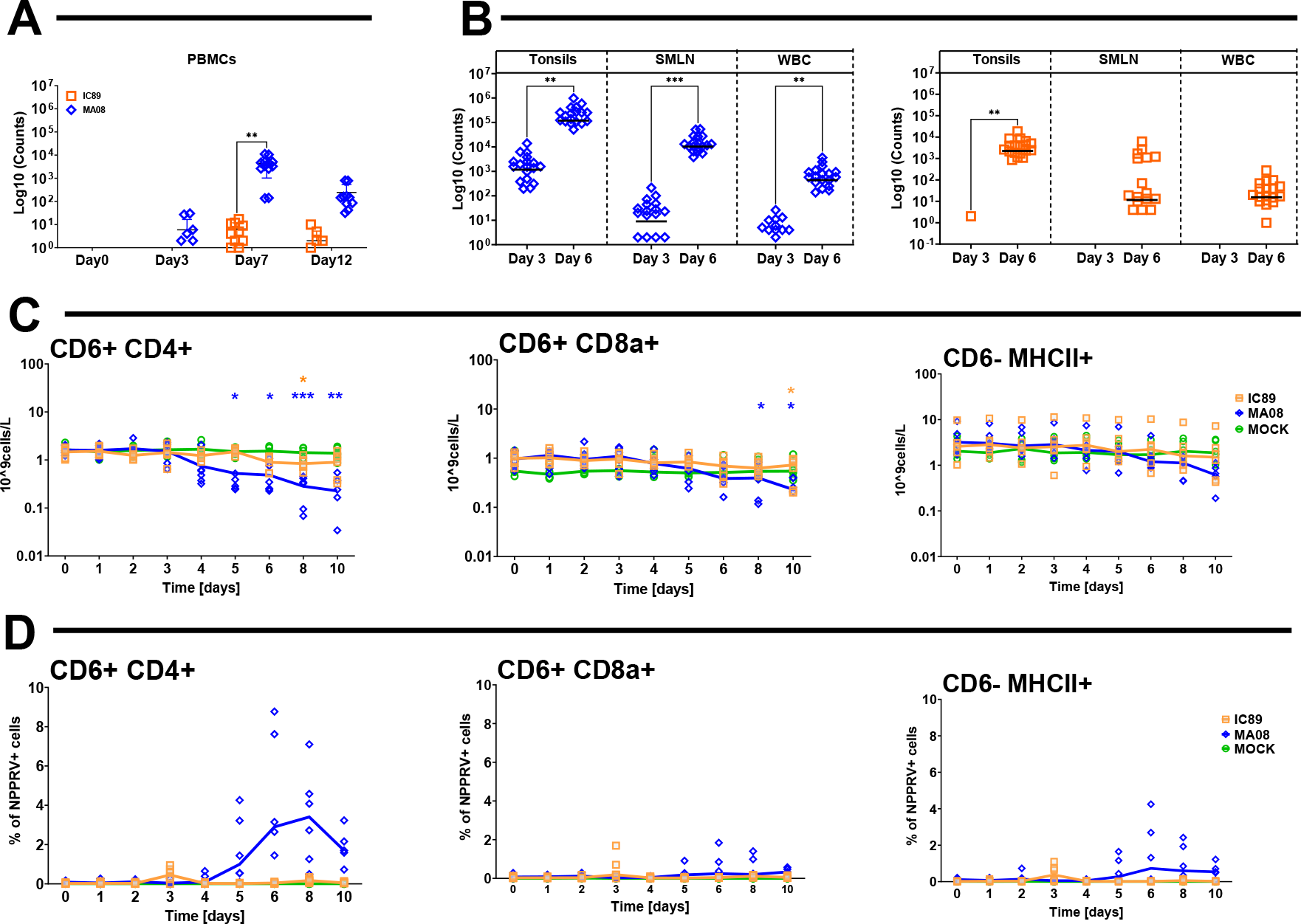
Impact of infection and detection of viral genes and nucleoproteins in different organs and cell compartments. In **A** and **B**, the number of reads aligned to the viral genome in raw transcriptome data from PBMCs (A), tonsils, superior mesenteric lymph node and white blood cells (B) is shown. In **C**, the absolute numbers of CD4 T cells (CD4+CD6+) CD8a T cells (CD6+CD8a+) and antigen-presenting cells (CD6-MHCII+) in circulating PBMCs is shown. In **D,** percentages of viral nucleoprotein in the PBMC populations are shown. In C, p-values were calculated by comparing group means of infected with MOCK-inoculated animals. *P-values < 0.05; **P-values < 0.01; ***P-values < 0.001.

Lymphocyte subpopulation dynamics monitored by flow cytometry revealed a significant depletion of CD4 T cells (CD6^+^CD4^+^) starting at 5 dpi with the MAO8. This was not observed following infection with the IC89 strain. The CD8 T cells only showed mild depletion at the end of the observation period, which appeared to be similar with both PPRV strains. CD6-MHCII cells were not affected (**Fig 2C**). PPRV nucleoprotein expression was predominantly found in the CD4 T cells starting at 5 dpi, specifically during MA08 infection (**Fig 2D**).

### PPRV strain-dependent innate immunity transcriptional profiling

#### PBMCs and WBCs

PBMCs and white blood cells (WBCs) that were collected in the two experiments showed similar trends on the modulation of innate immunity with a clear discrimination between the two viral strains (**Fig 3A-B**). At 3 dpi, the initiation of an interferon type I (IFN-I) response (M127) was only found in the goats infected with the MA08 strain (e.g. Blood transcriptional modules “BTM” M150, **Fig 3A**). Nevertheless, this response appeared to be more prominent at the later time points (6 and 7 dpi) with the IC89 strain, which induced more IFN-I BTM such as M111.1, M75 and M127. At 12 dpi, the IFN-I BTM returned to steady state with both groups (**Fig 3A**).

**Fig 3.**
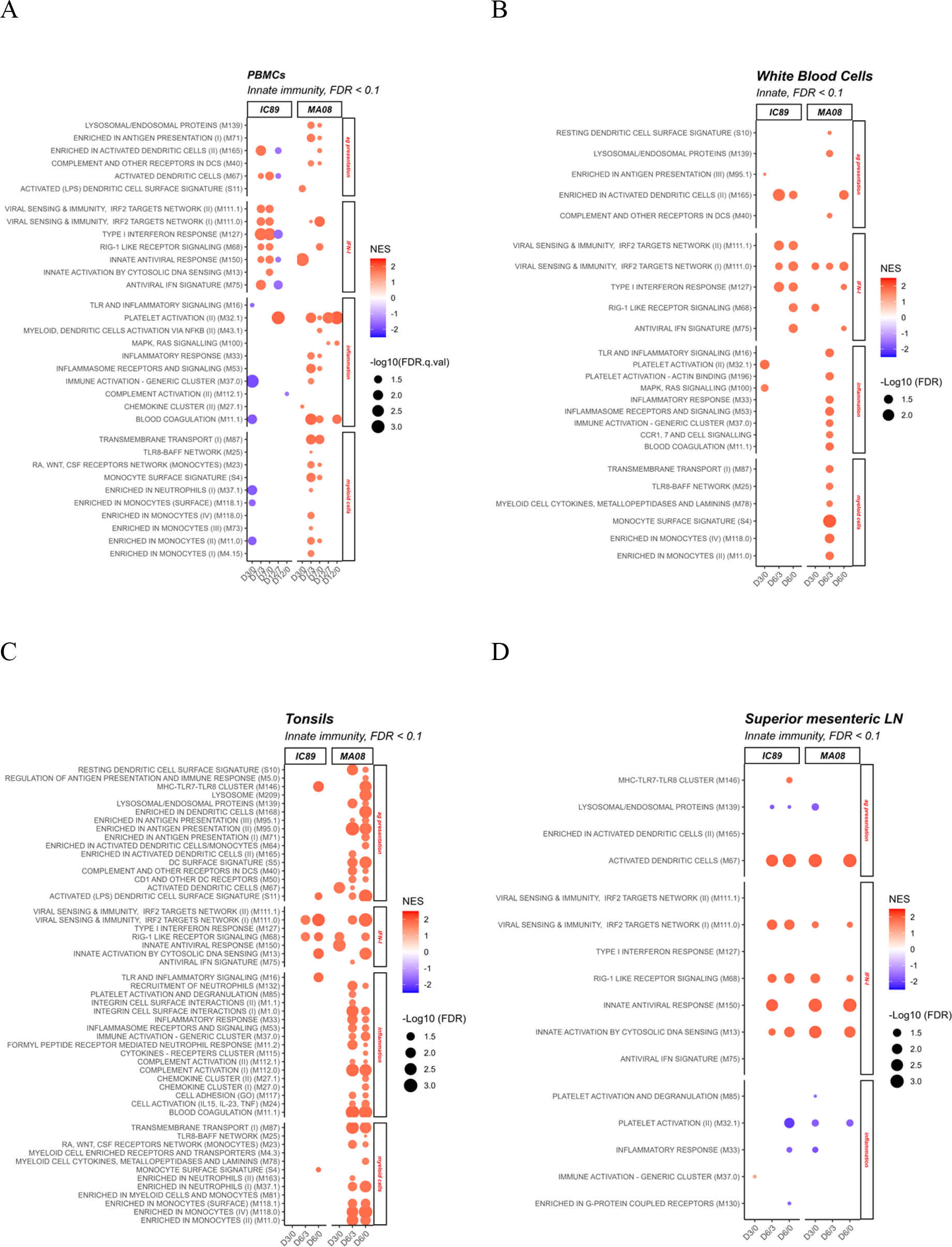
Innate-immunity related BTM perturbations induced by IC89 and MA08 PPRV strains. These plots show the induction (red) or downregulation (blue) of innate immunity BTM expression in PBMCs (**A**), tonsils (**B**), superior mesenteric lymph nodes (**C**) and white blood cells (**D**) at 3- and 6-days post-infection. The intensity of the colours reflects the Normalised Enrichment Scores (NES) determined after D0 to D3 (D3/0), D3 to D7 (D7/3), D0 to D7 (D7/0), D12 to D7 (D12/7) and D0 to D12 (D12/0) comparisons for PBMCs, and D0 to D3 (D3/0), D3 to D6 (D6/3) and D0 to D6 (D6/0) comparisons for other compartments. The size of the circles represents the FDR value. Families to which the significantly regulated BTMs correspond are shown on the right of the plots. D0 in these comparisons represents Mock- inoculated animals sacrificed 3 days after infections.

Analyses of the BTM related to inflammation and myeloid cells revealed a major difference in the host response between both PPRV strains (**Fig 3A**). Inflammatory and myeloid cell-related BTMs (M33, M53, M37.0, M11.1, M87, M25, S4, M118.0, M11.0) showed activation in D7/3 (in PBMCs) and D6/3 (in WBCs) comparisons in MA08-infected animals only (**Fig 3A** and **B**, respectively). Interestingly, the D7/0 and D6/0 did not show such strong induction in these BTM (**Fig 3A** and **B**, respectively). To explain this apparent contradiction, we had a look at the normalized enrichment scores (NES) for inflammatory and myeloid cell BTM at day 3 dpi and found that most of them were negative but did not reach statistical significance. IC89 only induced platelet activation (M32.1,) at 7 dpi in the PBMC dataset and at 3 dpi in the WBC dataset (**Fig 3B)**. Furthermore, in PBMC dataset a downregulation of the inflammatory BTM M37.0 was observed (**Fig 3A)**. BTM related to the antigen presentation related modules were mainly induced in D7/3 (in PBMCs, list them; **Fig 3A**) and D6/3 (in WBCs, list them; **Fig 3 B**) comparisons. IC89 infection consistently activated the M165 module in PBMCs and WBCs (**Fig 3A** and **B**, respectively), whereas the M40 and M139 modules were consistently activated in MA08 infection (**Fig 3A and B**).

#### Tonsils

In the tonsils two modules linked to the IFN-I response (M68 and M150, **Fig 3C**) were activated early at 3 dpi, confirming the earlier induction of an IFN-I response in the MA08- infected animals. Despite this, the IFN-I response was comparable for the two PPRV strains. In contrast only with the MA08 strain, a significant activation of many BTMs related to antigen presentation, inflammation and myeloid cells was observed in both the D6/0 and D6/3 comparisons. For the inflammation, this included BTM related to inflammatory cytokines responses (M16, M33, M53, M37.0 and M24), chemoattraction of inflammatory cells (M132, M11.2, M27.0), complement activation (M112.0) and coagulation (M11.1, M85) (**Fig 3C**).

#### Mesenteric lymph node

In mesenteric lymph node, both infections mainly led to the induction of BTM-related to type I IFN response (M13, M150, M68, M111.0) and activation of the “activated dendritic cells” BTM M67 antigen-presenting module. These modules were activated 3 or 6 dpi in animals infected with MA08 or IC89, respectively (**Fig 3D**). Strikingly, neither inflammatory nor myeloid cell BTM response were found in this lymph node, suggesting that inflammation was restricted to the tonsillar mucosal barrier site at the time of sampling.

In conclusion, these results indicate that the IFN-I response is induced earlier by MA08 but does not appear to fundamentally differ from that induced by IC89. In contrast, a major inflammatory, myeloid cell and antigen presenting cell responses were only induced in the tonsils by the MA08 strain.

#### PPRV strain-dependent adaptive immunity transcriptional profiling

Adaptive immunity BTM were divided into “cell cycle”, “B cell” and “T cell” BTM. The first and major response of B- and T-cell responses during an immune response is clonal expansion of antigen-specific lymphocytes, which is mediated by cell proliferation. This response is visible in the cell cycle BTM.

#### PBMCs and WBCs

In the WBC and PBMC, cell cycle BTM were initially downregulated at 3 and 6/7 dpi, followed by an upregulation at 12 dpi. While the downregulation of the cell cycle BTM was similar with the two PPRV strains, the upregulation at 12 dpi appeared more prominent with the MA08 strain (**Fig 4A, B**). For B- and T-cell BTM we observed only a very few downregulated BTM in the blood compartment.

**Fig 4.**
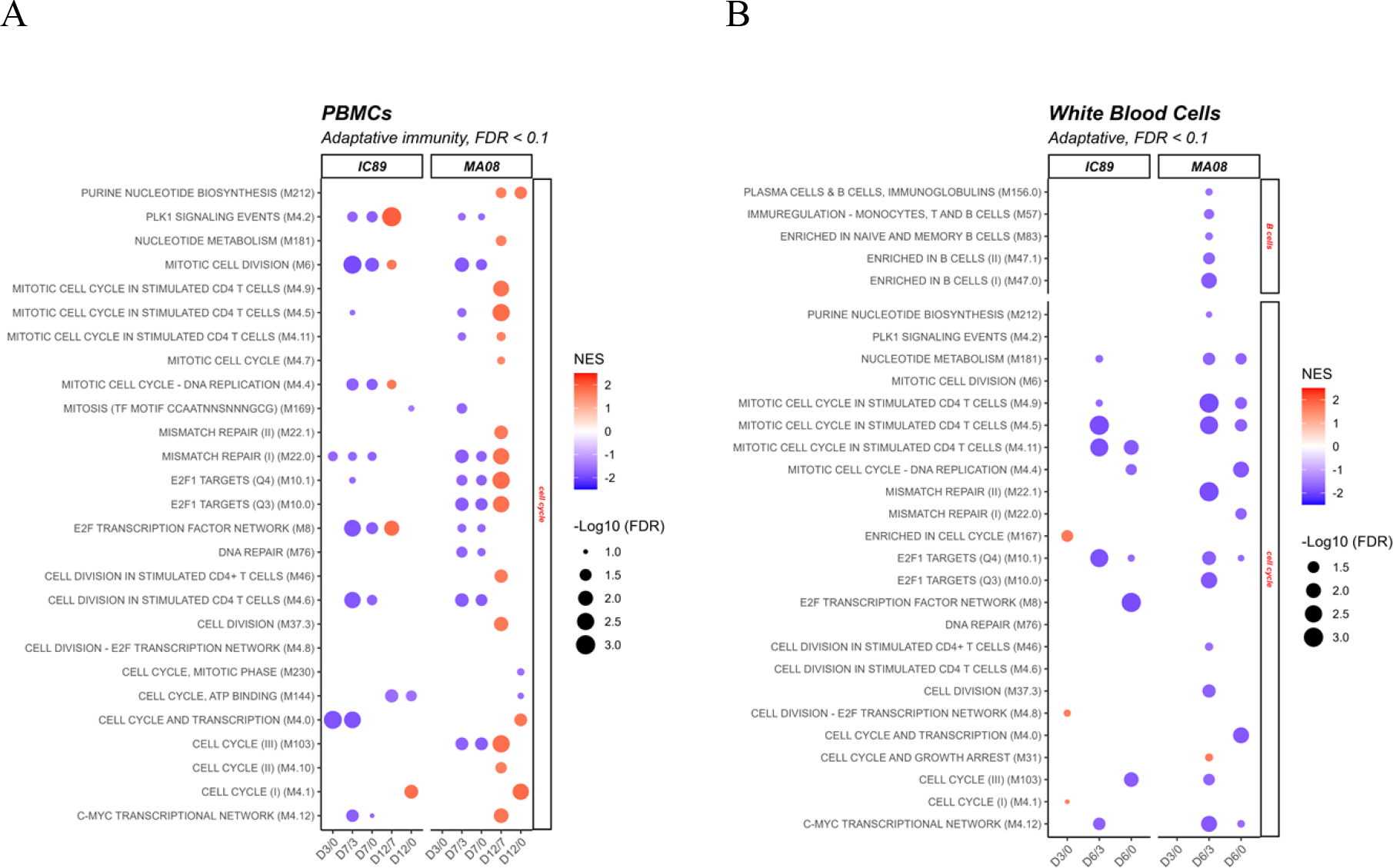
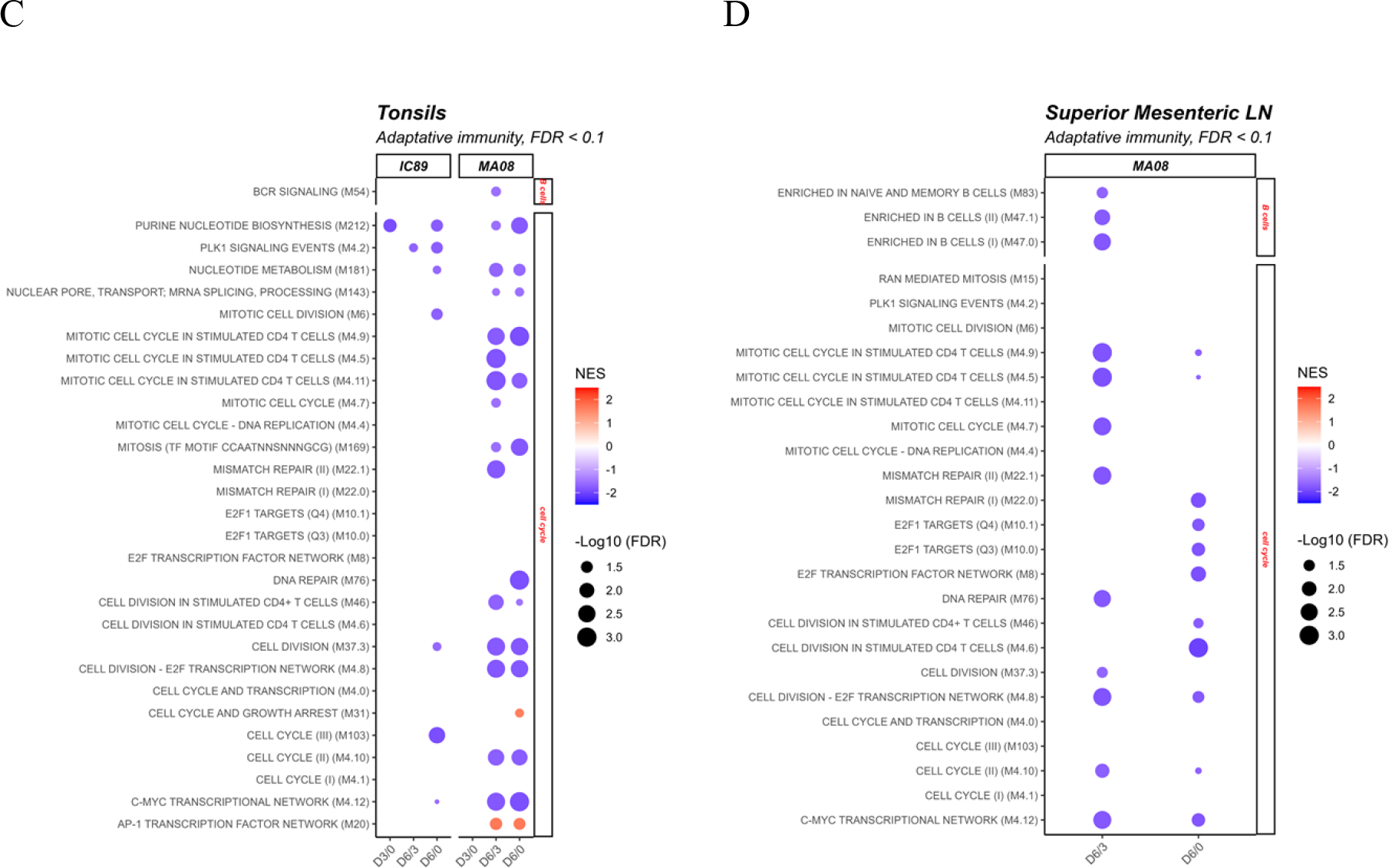
Adaptative-immunity related BTM perturbations induced by IC89 and MA08 PPRV strains. These plots show the induction (red) or downregulation (blue) of adaptative immunity BTM expression in PBMCs (**A**), white blood cells (**B**), tonsils (**C**) and superior mesenteric lymph nodes (**D**) at 3- and 6 days post infection. The intensity of the colours reflects the Normalised Enrichment Scores (NES) determined after D0 to D3 (D3/0), D3 to D7 (D7/3), D0 to D7 (D7/0), D12 to D7 (D12/7) and D0 to D12 (D12/0) comparisons for PBMCs, and D0 to D3 (D3/0), D3 to D6 (D6/3) and D0 to D6 (D6/0) comparisons for other compartments. The size of the circles represents the False Discovery Rate (FDR) value. Families to which the significantly regulated BTMs correspond are shown on the right of the plots.

#### Tonsils and mesenteric lymph nodes

In the tonsils and the mesenteric lymph nodes we observed a downregulation of cell cycle BTM at 6 dpi, which was much more prominent with the MA08 strain (**Fig 4C, D**). In lymph nodes we found no significant modulations with the IC89 strain (**Fig 4CD**). With respect to B- and T-cell BTM, similar to the blood, very few B cell BTM were downregulated in the D6/3 comparison in the lymph node following infection with MA08 (**Fig 4C, D**).

In conclusion, the transcriptional profiles of adaptive immunity indicate a significant suppression of proliferation in the lymphoid tissue during the first week following infection with the MA08 strain only. This inhibition of proliferation is also observed in the blood cells and is only reverted at 12 dpi.

### Proteome analysis confirmed the induction of a stronger innate immune response during infection with the highly virulent strain

Proteomics data from PBMCs collected at 0 and 6 dpi confirmed that infections by both strains mainly induced the synthesis of IFN-I related proteins, but in a more complex manner for MA08 infection. In addition, MA08 infection upregulated the synthesis of several large subunit ribosome proteins (RPL), inflammation-related proteins (S100A8, S100A9, S100A12, AZU1) and proteins that enhance replication of RNA viruses in a direct or indirect manner (ADAR1, PSFM1). Also, unnamed proteins such as na.18, na.86, LOC102179380 and 102173185 appeared to be upregulated and to play a role during MA08 infection.

Interestingly, MA08 infection led to a decrease in FLI-1 and na.18 expression, while MAVS, ACLS4, TJP2 and CCDC137 expression was decreased upon IC89 infection (**Fig 5A-B**).

**Fig 5.**
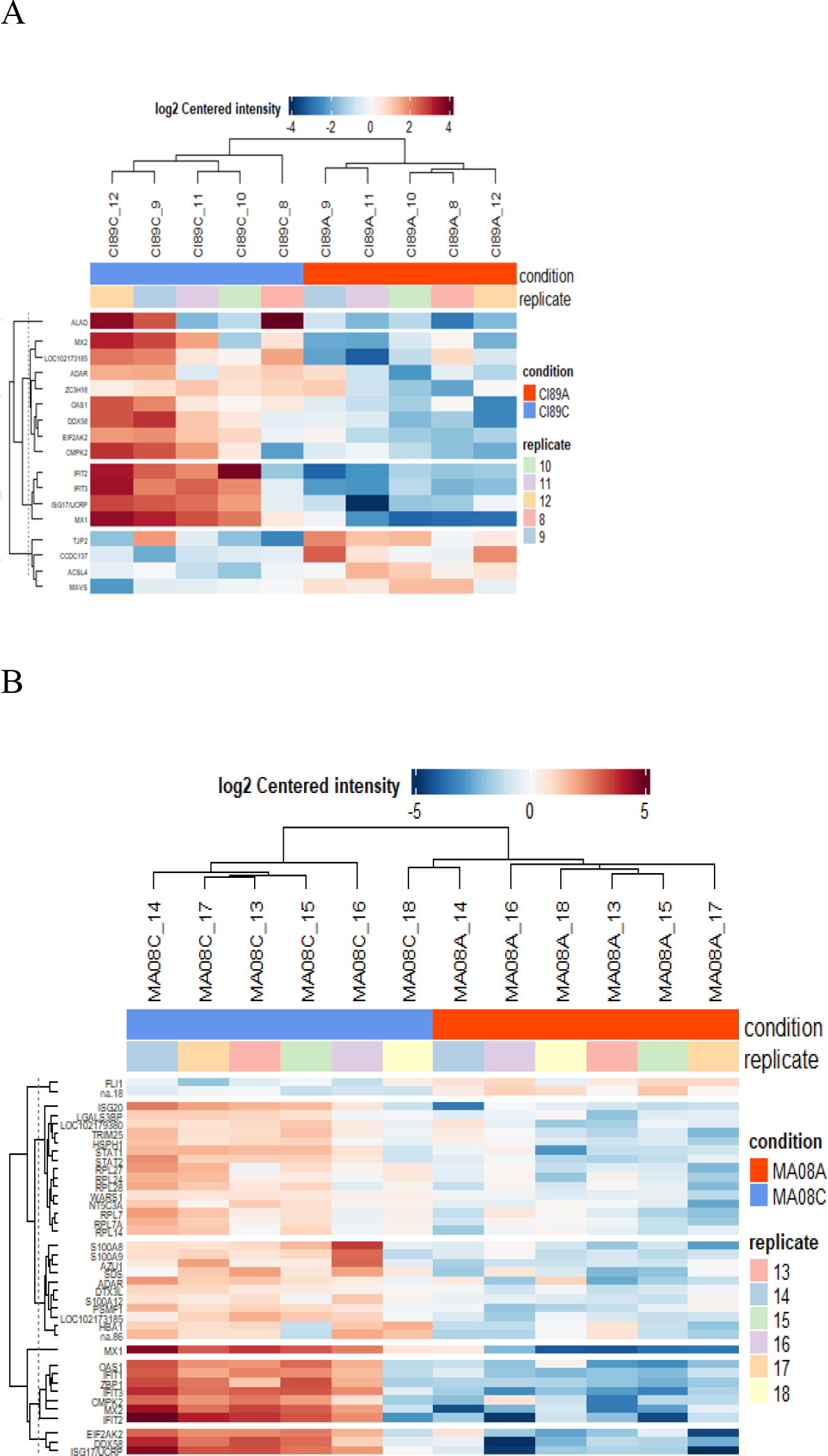
Proteome regulation analysis during PPRV Infection. Differentially Expressed Proteins (DEPs) statistically different between 0 (IC89A, MA08A) and 6 days (IC89C, MA08C) after infection with PPRV IC89 (**A**) or MA08 (**B**) are represented in these heatmaps. Each animal (replicate) within conditions is represented with squares of different colours. Clustering was performed on rows (expressed proteins) and columns (different replicates).

Network prediction showed that these ribosomal genes are linked to ISG15 expression (**S2 Fig**).

### PPRV infection of tonsillar epithelium by MA08 is associated with tonsillar inflammation

The transcriptomic analyses indicated severe inflammation and myeloid infiltration in the tonsils following MA08 infection. Considering that this effect was not observed in the mesenteric lymph node (MLN) at the time of sampling, we postulated that PPRV infection in the tonsils could induce barrier defects that promote innate immune responses induced by bacteria present in the tonsillar crypts. Indeed, immunohistochemistry (IHC) for viral nucleoprotein demonstrated high level of infection in the tonsillar epithelium in MA08 infected goats at 6 dpi, with a prominent positivity within the underlying lymphoid tissue.

Moreover, the tonsillar epithelium showed histologically multifocal ulcerations, which were associated with haemorrhages and fibrin exudation, as well as abundant leukocytes, predominantly neutrophils, admixed with bacteria in the lumen of the crypts (**Fig 6**). It is conceivable that the PPRV infection could cause the observed inflammatory responses directly or indirectly by disturbing the epithelial barrier function. In contrast, this inflammatory response was absent in MLN, despite high viral loads in MA08-inoculated goats at 6 dpi (**Fig 7**).

**Fig 6.**
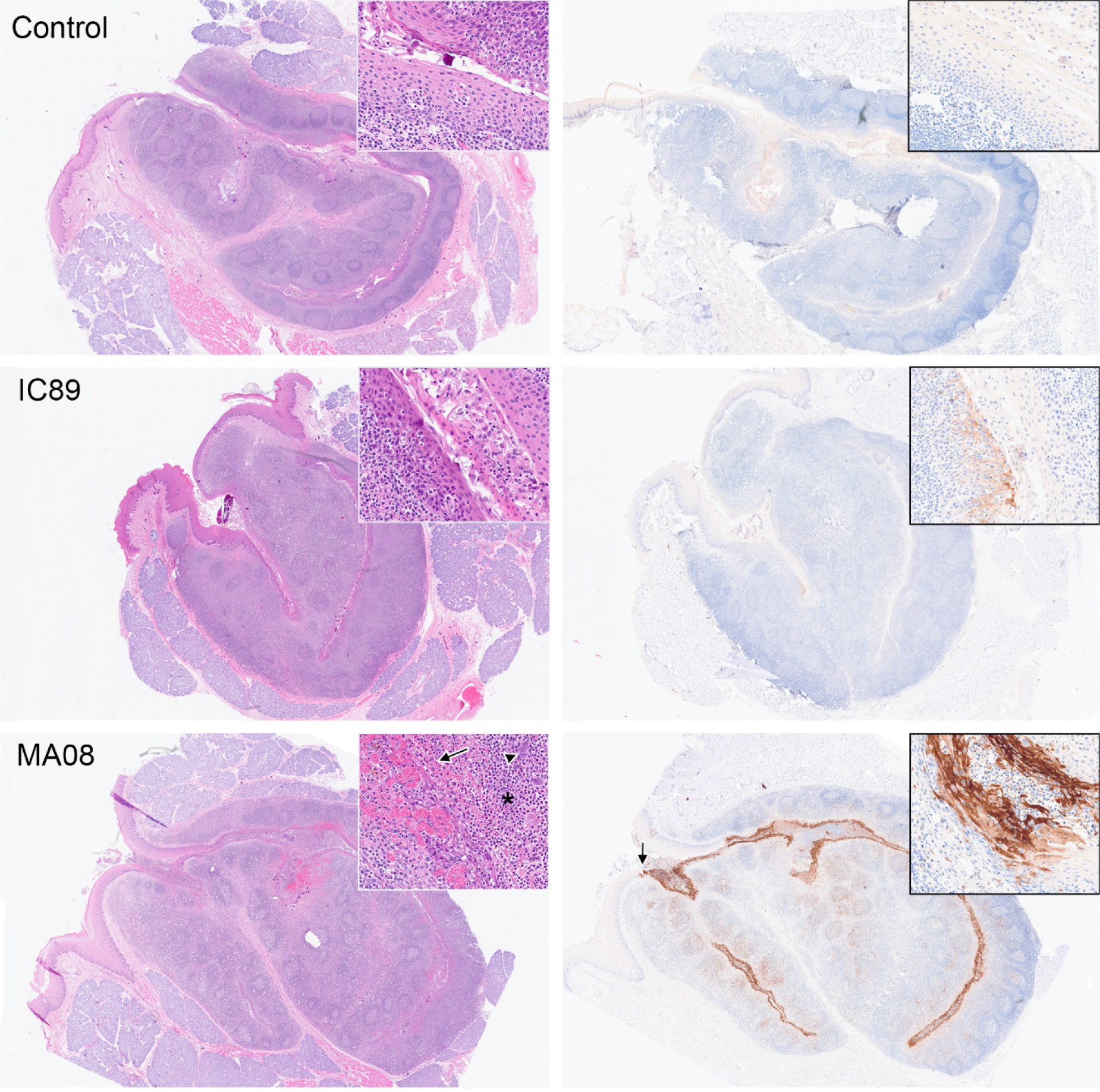
Panel showing histological and immunohistochemistry (IHC) pictures of tonsils from one control, one IC89-inoculated and one MA08-inoculated goat at 6 dpi, representative of each experimental group. Insets show higher magnification details of the tonsillar epithelium. Left: Haematoxylin eosin (HE). Both control and IC89-infected goat show multifocal infiltration of leukocytes within the tonsillar epithelium, and rather low numbers of leukocytes and some bacteria in the lumen of the crypts. MA08-infected goat shows multifocal attenuation and ulceration of the tonsillar epithelium associated with areas of haemorrhage (arrow), some fibrin exudation and with abundant neutrophils (asterisk) and bacteria (arrowhead) in the tonsillar lumen. Right: IHC for PPRV protein N. Very strong and diffuse intracytoplasmic staining (brown) is observed in the tonsillar epithelium of the MA08 infected goat, with a very sharp demarcation with the oropharyngeal epithelium (arrow). A prominent IHC PRRV positivity is also observed in the underlying follicular and parafollicular tonsillar tissue. In contrast, a weak multifocal positivity is observed in the tonsillar epithelium and underlying lymphoid tissue in the IC89 infected goat.

**Fig 7.**
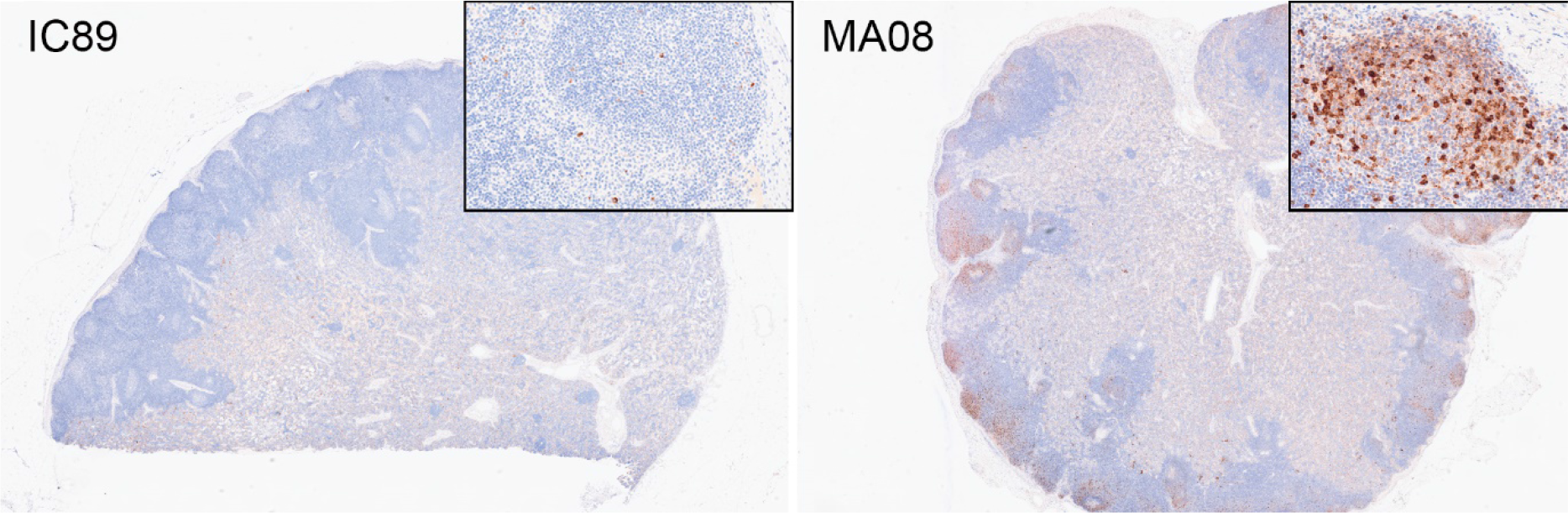
Immunohistochemistry for PPRV in mesenteric lymph nodes from one IC89- inoculated and one MA08-inoculated goat at 6 dpi, representative of both experimental groups. IC89 showed only a mild scattered intracytoplasmic brown staining within the lymph node parenchyma, while MA08 showed abundant positivity, which seems to be more prominent but not restricted to the lymphoid follicles.

The IHC intensity and distribution within the lymphoid tissue in MA08-inoculated goats was similar in the tonsils and the MLN. Thus, at 6 dpi there was a moderate to marked positivity in both follicular and parafollicular areas. In the tonsil, the strongest positivity was seen within the crypt epithelium of the tonsils. On the contrary, in IC89-innoculated goats at 6 dpi showed only few scattered stained individual cells within the lymphoid tissue, with few small, multifocal positively stained areas in the tonsillar crypt epithelium.

## Discussion

To understand the biological basis of strain-dependent PPRV virulence, we have used an experimental model, based on the infection of Saanen goats or their PBMC with PPRV strains of different virulence[22,7]. In goats, the MA08 strain induced higher clinical scores associated with more prominent lymphocyte depletion, and lesions particularly affecting mucosal and lymphoid tissues. As in vitro model we have used PBMCs from Saanen goats that were stimulated with mitogen and then infected with either IC89 or MA08. This revealed that viral replication prevents in vitro lymphocyte proliferation and promotes cell death [23]. The present work aims to understand early events in viral immunopathogenesis related to these virulence-dependent features. To this end, we focussed on selected lymphoid tissues either in direct contact with mucosal surfaces (tonsils) or in lymph nodes draining mucosal tissue (MLN).

The nasopharyngeal epithelium and tonsillar tissues represent the likely initial entry site for PPRV where the virus first replicates in epithelial cells followed by antigen-presenting cells [24,25]. Thereafter, infected antigen-presenting cells will reach lymph nodes draining the inoculation site [26]. Our results are in line with this, showing that tonsils have the highest viral load in terms of viral RNA, starting 3 dpi for the highly virulent MA08 and 6 dpi for the low virulent IC89 strain. Accordingly, following MA08 infection, innate BTM including inflammation, myeloid cells and antigen-presentation related modules are strongly activated in tonsils. However, it is quite remarkable that this inflammatory response was completely absent in MLN, despite high viral loads and a prominent presence of PPRV protein N at 6 dpi. A possible explanation is that a massive infection of the tonsillar epithelium by PPRV associated with the observed tonsillar epithelial damage leads to barrier defects, promoting inflammatory responses. Activation of inflammation-related modules may either be directly linked to the strong presence of PPRV protein N, which plays an important pro-inflammatory role by modulating inflammasome assembly [18]. Alternatively, the detected inflammation may be induced by the bacteria, which are often being seen in the lumen of the crypts, being triggered by the breach in the tonsillar epithelium and therefore loss of this important anatomical and immunological barrier. A combination of both phenomena is also possible. In contrast, the MLN does not constitute a direct mucosal barrier. It can however not be totally ruled out that a similar inflammatory response and epithelial damage may occur in the intestine. Additional experiments would be needed to resolve this question. Animals infected with the less virulent strain did not show damage in the tonsillar epithelium nor showed such massive infection or activation, which suggests that the anatomical and immunological barrier operates at the site of entry in the tonsillar crypts. It must be noted that in formalin-fixed tissues from natural measles cases, the majority of measles virus-infected cells in epithelium were the intraepithelial CD11c+ myeloid cells[27]. Furthermore, it has been speculated that resident myeloid cells in the respiratory mucosa can transport PPRV^21^. Future experiments are needed to understand the role of epithelial and myeloid cells in the inflammatory responses associated with highly virulent PPRV only. For example, differences in viral tropism for specific cell types might play a role in virulence.

Besides the induction of strong inflammatory responses in tonsils, our transcriptomic data also revealed an early activation of IFN-I responses at 3 dpi with the highly virulent MA08 strain. Nevertheless, at 6 dpi the IFN-I related BTM were expressed at levels comparable to those induced by IC89. This indicates that IFN-I response during PPRV is not involved in pathogenic effects such as severe inflammation and lymphoid depletion.

Our results also demonstrate another prominent effect of virulent PPRV strains on the lymphoid tissue, which was the downregulation of the cell cycle BTM in the first week of infection. Proliferative responses in lymphoid tissue are of central importance because they indicate the initiation of adaptive immune responses and correlate well with clonal expansion and the induction of potent T cell responses. Based on this, cell cycle BTM have been demonstrated to represent a correlate of T cell responses[28–31], and were found to be induced in the first week of a virus infection[31,32]. Based on the opposite effect induced by PPRV and observations from other morbilliviruses[11–16] we propose that the suppression of these responses is part of the pathogenesis of virulent PPRV infection. The cell cycle arrest may be linked to the significant reduction in circulating CD4+ T lymphocytes observed in goats at 6 dpi with MA08 strain. One possible pathway for inducing this immunosuppression may involve PPRV-infected dendritic cells, which have been shown to suppress the proliferative T-cell response[15]. This may be in line with the observed depletion of CD4+ T lymphocytes reported by others [33]. In addition, we now also show that CD4+ T cells are a main target of PPRV infection in the blood circulation during virulent PPR, and this could contribute to immunosuppression.

It is important to note that goats can overcome the immunosuppression induced by virulent PPRV. On one side MA08 did not prevent the induction of neutralizing antibodies, which were first detected at around 7 dpi[7] . On the other hand, cell cycle BTM were induced at 12 dpi. This suggest that neutralizing antibodies are essential and not sufficient in controlling this infection and the importance of the induction of PPRV specific T-cell responses. Evidently, the general health of the animal at the time of infection will influence the efficacy of its immune response.

Our analysis of the proteome of PBMCs confirmed that infections are associated with the innate antiviral response in both low and high virulent PPRV strains. It has also been shown that after vaccinating goats with the Sungri 96 PPRV vaccine, a majority of proteins associated with the antiviral interferon response were synthesized [34]. With MA08 infection, we identified ribosomal proteins that may be linked to ISG15 expression in line with other studies [35]. Nevertheless, most ribosomal protein-virus interactions are essential for virus translation and replication, while a minority activate immune responses against the virus [36,37]. Therefore, we could hypothesize that these proteins are involved in the elevated replication of MA08. Indeed, differentially acetylated ribosomal proteins were found following PPRV infection of Vero cells [38]. The inflammation observed in MA08-infected animals may also be linked to protein expression, as a core set of inflammatory proteins was expressed in PBMCs. In this context, this strain is able to use all the mechanisms described elsewhere that are involved in PPRV virulence[39,17,19,40–43].

In conclusion, comparison of low and high virulent PPRV strains using a multiomics approach has identified crucial early innate and adaptive immunopathological processes associated with severe inflammatory disease and ineffective control of the virus by adaptive immune responses. This information is required to understand strain- and host-dependent differences in pathogenesis of PPRV, which are relevant to PPRV surveillance and control strategies

## Methods

### Ethics statement

Animal experimentations were carried out in the high containment facility of the Institute of Virology and Immunology (IVI, Mittelhäusern), in accordance with the Swiss animal protection law (TSchG SR 455; TSchV SR 455.1; TVV SR 455.163). The committee on animal experiments of the canton of Bern, Switzerland, reviewed the experiments, and the cantonal veterinary authority approved the study under the authorization number BE16/2020.

### Animal experiments

Samples used in this study were collected during two different animal experiments. The first animal experimentation was previously described [7]. Briefly, twenty-four Saanen goats aged between two and twelve years were randomly distributed into four groups (mock-inoculated, vaccinated, IC89-infected and MA08-infected) of six animals. Clinical signs as well as the gross pathology and seroconversion of the animals were described. Blood samples were collected in all groups at days 0, 3, 7 and 12 post infection to study the modulation of RNA over the time course of the experiment (Fig 1.A). Based on data obtained in this first experiment (see below), we decided to further investigate transcriptional changes in different lymphoid organs at the early stage of infection with PPRV strains of different virulence. For this purpose, eighteen adults Saanen goats were randomly distributed into three groups (mock-inoculated, IC89-infected and MA08-infected) of six animals. Tonsils, superior mesenteric lymph nodes (MLN) and white blood cells (WBCs) were collected at three and six days after inoculation. For each sampling day, three animals were sacrificed for organ collection. Clinical signs of disease were monitored throughout the experiment (Fig 1.B).

### Flow cytometry

Peripheral blood mononuclear cells (PBMC) collected from the first experiment and prepared as previously described[23] were analyzed to quantify T helper cells, defined as CD4^+^CD6^+^, cytolytic T cells defined as CD8^+^CD6^+^ and antigen presenting cells, defined as CD6^-^MHCII^+^. The latter contained mainly B cells, monocytes and dendritic cells. Primary antibodies used were anti-CD6 (IgG1, clone BAQ91A), anti-CD4 (IgG2a, clone GC1A), anti-CD8α (IgG2a, clone 7C2B) and anti-MHCII (IgG2a, clone TH16A), all purchased from the Monoclonal Antibody Center (Washington State University, USA). Secondary antibodies conjugated with PE-Cy7 (anti IgG1) and Alexa Fluor 647 (anti IgG2a) were purchased from ThermoFisher Scientific. For the intracellular staining of the virus, cells were fixed in 4% formalin solution (PFA) and permeabilised in 1X PBS supplemented with 0.1% of Saponin. Fixed and permeabilised cells were stained with FITC-conjugated NPPRV[44] antibodies (clone 38-4, CIRAD, Montpellier, France) diluted (1:100) in PBS supplemented with 0.3% of Saponin.

### Histopathology and immunohistochemistry

Tonsil and MLN samples were placed in 10% buffered formalin, routinely processed for histology, sectioned at 3 μm and stained with haematoxylin and eosin (H&E). For immunohistochemistry (IHC) 3 μm formalin-fixed, paraffin-embedded tissue sections were mounted on positively charged slides (Color Frosted Plus, Biosystems, Muttenz, Switzerland), dried for 35 min at 60°C and subsequently dewaxed, pretreated and stained on Bond-III immunostainers (Leica Biosystems, Melbourne, Australia). After dewaxing (Bond Dewax solution; Leica Biosystems), slides were subject to a heat induced epitope retrieval step, using a Tris-EDTA based buffer (Bond Epitope Retrieval 2; pH 9) for 10 min at 95°C to 100°C. To reduce non-specific binding of primary antibodies a protein block solution was applied for 10 min at room temperature, as for all following steps. Then the slides were incubated with a primary mouse monoclonal, antibody against the PPRV nucleocapsid, clone 38-4[44], at 1:100 for 15 min. All further steps were performed using reagents of the Bond Polymer Refine Detection Kit (Leica Biosystems) as follows: Endogenous peroxidase was blocked for 5 min, then a rabbit-anti-mouse secondary antibody was applied (8 min), followed by a peroxidase-labelled polymer (8 min). Both these reagents were supplemented with 2% dog serum to block non-specific binding (LabForce, Nunningen, Switzerland). Finally, slides were developed in 3,3’-diaminobenzidine / H_2_O_2_ (10 min), counterstained with haematoxylin, and mounted. In negative controls the primary antibody was replaced with wash buffer. Known positive controls were stained in parallel with each series.

### RNA Sequencing

Total RNA was extracted from caprine PBMCs, WBCs and organs using the protocol described in Eloiflin et al. [23]. Total RNA quality and concentration were measured, and the library prepared at the Next Generation Sequencing platform of the University of Bern, Switzerland. Only samples with RNA Quality Number (RQN) score ≥ 8 were sequenced on a NovaSeq 6000 instrument using an SP flow cell. Raw sequencing reads were aligned on either *Capra hircus* ARS1 genome (GCA_001704415.1) or PPRV genome (GCA_000866445.1 ViralProj15499) using STAR 2.7.3a [45]. Before mapping step, STAR index was generated from FASTA and annotation GTF files of selected references with the following parameters: star --runThreadN –runMode --genomeDir --genomeFastaFiles --sjdbGTFfile --sjdbOverhang. Read mapping was then performed on the generated index using the following parameters: STAR --genomeDir –runThreadN –readFilesCommand --readFilesIn --outFileNamePrefix -- outSAMtype BAM --outSAMunmapped --peOverlapNbasesMin --outSAMattributes.

Quantification of gene expression was performed using FeatureCounts v2.0.1 [46]. Reads were assigned to the reference annotations at the exon level and counts were summarized at the gene level, using default parameters (-t exon -g gene_id), corresponding to unambiguously assigning uniquely aligned paired-end (-p) and multi-mapped reads (-M). Then, the obtained counts table was used for differential expression analysis between not-infected (Mock-inoculated) and infected animals at 3 and 6 days post-infection with DESeq2 v1.36.0 [47] on R software v4.2.2. Before running the differential expression analysis, clustering of replicates was verified by running a principal component analysis with plotPCA function of the DESeq2 package.

### Gene set enrichment analysis

Differentially expressed gene (DEG) lists obtained after comparisons were ranked to enable gene set enrichment analysis (GSEA) using GSEA v4.3.0 desktop application [48,49]. This ranking assigns each gene a unique rank value based on its expression value (log2 fold changes) and its adjusted p-value. Thus, each gene’s rank is calculated using the following formula: *RNK = - Log (adjusted p-value) * SIGN*; where SIGN is “-1” when the gene is down-regulated and “+1” when it is up-regulated. These ranked files were then loaded into the software. Blood transcriptional modules (BTM) adapted to *Capra hircus* as well as a chip annotations file that lists each identifier and its matching HGNC gene symbol used in BTM [23] were also loaded in the software and used first to run the Chip2Chip mapping tools with default parameters. This step converts gene sets to the format required for the chip platform and allows analysis of the dataset without reducing the probe sets to gene symbols. Ranked files and BTMs mapped to the chip platform were used to run the GSEAPreranked mode with the following parameters: required fields: -- Number of permutations 1000; --Collapse/Remap to gene symbols No_Collapse and basic fields: --Enrichment statistic weighted; --Max size: exclude larger sets 500; --Min size: exclude smaller sets 5. All other fields were used with the default parameters. BTM were classified in families relating to “antigen presentation”, “IFN type I”, “myeloid cells”, “B cells”, “T cells” and “cell cycle” as previously described[23]. The enrichment score (ES) and adjusted p-values for the BTM were visualized with ggplot2 in R software.

### Proteomics

During the first animal experiment and after extraction, PBMCs were also collected in DIGE buffer (7 M urea, 2 M thiourea, 4% 3-[(3-cholamidopropyl)-dimethylammonio]- 1propanesulfonate (CHAPS) and 30 mM TrisHCL; pH 8). These samples were prepared and analysed by mass spectrometry following the protocol described in Eloiflin et al.[23]. Data obtained after mass spectrometry analysis were used to identify differentially expressed proteins in our samples. A heatmap clustering analysis of the log2 Intensity values for the most significantly expressed proteins between conditions was conducted using Pheatmap library on R software v4.2.2.

### Statistical analysis

Statistical tests were carried out with Graphpad Prism 9.4.1 (GraphPad Software, California, USA). A mixed effects model (REML) was used with the Geisser-Greenhouse correction. In this model the measures recorded over time for each of the animals were considered as matched values. P-values between the mean of groups (MOCK, IC89 or MA08-infected animals) were calculated using Tukey’s multiple comparison test. Individual’s variances were computed for each comparison. Asterisks on the graphs highlight statistical differences between the comparisons. *P-values < 0.05; **P-values < 0.01; ***P-values < 0.001; ****P- values < 0.0001.

## Data accessibility

Sequencing data have been deposited in NCBI SRA (BioProject ID SUB14393662).

## Supporting information

**S1 Fig.** RT-qPCR on organs collected during the experiment.

**S2 Fig.** Protein-protein interaction networks generated 6 days after infection of Saanen goats with PPRV strains IC89 (A) and MA08 (B).

## Acknowledgments

The authors would like to thank the staffs of the UMR ASTRE (CIRAD) and the IVI (BERN) for their help with this project.

## Author Contributions

Conceptualization: Roger-Junior Eloiflin, Philippe Totte, Obdulio García- Nicolás, Artur Summerfield, Arnaud Bataille.

Data Curation: Roger-Junior Eloiflin, Artur Summerfield

Formal Analysis: Roger-Junior Eloiflin, Llorenç Grau-Roma, Artur Summerfield Funding Acquisition: Arnaud Bataille, Artur Summerfield

Investigation: Roger-Junior Eloiflin, Vincent Lasserre, Sylvie Python, Obdulio García- Nicolás, Llorenç Grau-Roma

Methodology: Roger-Junior Eloiflin, Stephanie Talker, Obdulio García- Nicolás, Llorenç Grau-Roma, Philippe Totte, Artur Summerfield

Supervision: Arnaud Bataille, Artur Summerfield Validation: Philippe Totte, Artur Summerfield Visualization: Roger-Junior Eloiflin, Llorenç Grau-Roma Writing – Original Draft Preparation: Roger-Junior Eloiflin

Writing – review & editing: Roger-Junior Eloiflin, Llorenç Grau-Roma, Vincent Lasserre, Sylvie Python, Stephanie Talker, Philippe Totte, Obdulio García- Nicolás, Artur Summerfield, Arnaud Bataille.

## Notes

### Competing Interest Statement

The authors have declared no competing interest.

## References

1. Parida S, Muniraju M, Mahapatra M, Muthuchelvan D, Buczkowski H et al. (2015) Peste des petits ruminants. Veterinary microbiology 181 (1-2): 90–106. Available: https://pubmed.ncbi.nlm.nih.gov/26443889/.

2. Njeumi F, Bailey D, Soula JJ, Diop B, Tekola BG (2020) Eradicating the Scourge of Peste Des Petits Ruminants from the World. Viruses 12 (3).

3. Fine AE, Pruvot M, Benfield CTO, Caron A, Cattoli G et al. (2020) Eradication of Peste des Petits Ruminants Virus and the Wildlife-Livestock Interface. Frontiers in veterinary science 7: 50.

4. Baron MD, Diallo A, Lancelot R, Libeau G (2016) Peste des Petits Ruminants Virus. Advances in virus research 95: 1–42.

5. Baron MD, Diop B, Njeumi F, Willett BJ, Bailey D (2017) Future research to underpin successful peste des petits ruminants virus (PPRV) eradication. The Journal of general virology 98 (11): 2635–2644.

6. Couacy-Hymann E, Bodjo C, Danho T, Libeau G, Diallo A (2007) Evaluation of the virulence of some strains of peste-des-petits-ruminants virus (PPRV) in experimentally infected West African dwarf goats. Veterinary journal (London, England : 1997) 173 (1): 178–183.

7. Eloiflin R-J, Grau-Roma L, Python S, Mehinagic K, Godel A et al. (2022) Comparative pathogenesis of peste des petits ruminants virus strains of difference virulence. Veterinary research 53 (1): 57.

8. 8. WOAH & FAO (2015) Global control and eradication of PPR. Available: https://www.fao.org/3/i4477e/i4477e.pdf. Accessed 12 December 2023.

9. 9. Albina E, Kwiatek O, Minet C, Lancelot R, Servan de Almeida R, et al. (2013) Peste des Petits Ruminants, the next eradicated animal disease. Veterinary microbiology 165 (1-2): 38–44.

10. Tatsuo H, Ono N, Yanagi Y (2001) Morbilliviruses use signaling lymphocyte activation molecules (CD150) as cellular receptors. Journal of virology 75 (13): 5842–5850.

11. Krakowka S, Cockerell G, Koestner A (1975) Effects of canine distemper virus infection on lymphoid function in vitro and in vivo. Infection and immunity 11 (5): 1069–1078.

12. McChesney MB, Altman A, Oldstone MB (1988) Suppression of T lymphocyte function by measles virus is due to cell cycle arrest in G1. Journal of immunology (Baltimore, Md. : 1950) 140 (4): 1269–1273.

13. Heaney J, Barrett T, Cosby SL (2002) Inhibition of in vitro leukocyte proliferation by morbilliviruses. Journal of virology 76 (7): 3579–3584.

14. Vries RD de, McQuaid S, van Amerongen G, Yüksel S, Verburgh RJ, et al. (2012) Measles immune suppression: lessons from the macaque model. PLoS pathogens 8 (8): e1002885.

15. Rodríguez-Martín D, García-García I, Martín V, Rojas JM, Sevilla N (2022) A Morbillivirus Infection Shifts DC Maturation Toward a Tolerogenic Phenotype to Suppress T Cell Activation. Journal of virology 96 (18): e0124022.

16. McChesney MB, Kehrl JH, Valsamakis A, Fauci AS, Oldstone MB (1987) Measles virus infection of B lymphocytes permits cellular activation but blocks progression through the cell cycle. Journal of virology 61 (11): 3441–3447.

17. Zhang R, Lin H, You Q, Zhang Z, Bai L et al. (2022) Peste des Petits Ruminants Virus Upregulates STING to Activate ATF6-Mediated Autophagy. Journal of virology 96 (20): e0137522.

18. Li L, Yang W, Ma X, Wu J, Qin X et al. (2022) Peste Des Petits Ruminants Virus N Protein Is a Critical Proinflammation Factor That Promotes MyD88 and NLRP3 Complex Assembly. Journal of virology 96 (10): e0030922. Available: https://pubmed.ncbi.nlm.nih.gov/35502911/.

19. Sanz Bernardo B, Goodbourn S, Baron MD (2017) Control of the induction of type I interferon by Peste des petits ruminants virus. PloS one 12 (5): e0177300.

20. Jin L, Li Y, Pu F, Wang H, Zhang D et al. (2021) Inhibiting pyrimidine biosynthesis impairs Peste des Petits Ruminants Virus replication through depletion of nucleoside pools and activation of cellular immunity. Veterinary microbiology 260: 109186.

21. 21. Li L, Li S, Han S, Li P, Du G, et al. (2023) Inhibition of caspase-1-dependent apoptosis suppresses peste des petits ruminants virus replication. Journal of veterinary science 24 (5).

22. Enchery F, Hamers C, Kwiatek O, Gaillardet D, Montange C et al. (2019) Development of a PPRV challenge model in goats and its use to assess the efficacy of a PPR vaccine. Vaccine 37 (12): 1667–1673.

23. Eloiflin R-J, Auray G, Python S, Rodrigues V, Seveno M et al. (2021) Identification of Differential Responses of Goat PBMCs to PPRV Virulence Using a Multi-Omics Approach. Frontiers in immunology 12: 745315.

24. Gautam S, Joshi C, Sharma AK, Singh KP, Gurav A et al. (2021) Virus distribution and early pathogenesis of highly pathogenic peste-des-petits-ruminants virus in experimentally infected goats. Microbial pathogenesis 161 (Pt A): 105232.

25. 25. Schmitz KS, Eblé PL, van Gennip RGP, Maris-Veldhuis MA, Vries RD de, et al. (2023) Pathogenesis of wild-type- and vaccine-based recombinant peste des petits ruminants virus (PPRV) expressing EGFP in experimentally infected domestic goats. The Journal of general virology 104 (2).

26. Pope RA, Parida S, Bailey D, Brownlie J, Barrett T et al. (2013) Early events following experimental infection with Peste-Des-Petits ruminants virus suggest immune cell targeting. PloS one 8 (2): e55830.

27. Allen IV, McQuaid S, Penalva R, Ludlow M, Duprex WP et al. (2018) Macrophages and Dendritic Cells Are the Predominant Cells Infected in Measles in Humans. mSphere 3 (3).

28. Li S, Rouphael N, Duraisingham S, Romero-Steiner S, Presnell S et al. (2014) Molecular signatures of antibody responses derived from a systems biology study of five human vaccines. Nature immunology 15 (2): 195–204.

29. 29. Qi Q, Cavanagh MM, Le Saux S, Wagar LE, Mackey S, et al. (2016) Defective T Memory Cell Differentiation after Varicella Zoster Vaccination in Older Individuals. PLoS pathogens 12 (10): e1005892.

30. Matthijs AMF, Auray G, Jakob V, García-Nicolás O, Braun RO et al. (2019) Systems Immunology Characterization of Novel Vaccine Formulations for Mycoplasma hyopneumoniae Bacterins. Frontiers in immunology 10: 1087.

31. Bocard LV, Kick AR, Hug C, Lischer HEL, Käser T et al. (2021) Systems Immunology Analyses Following Porcine Respiratory and Reproductive Syndrome Virus Infection and Vaccination. Frontiers in immunology 12: 779747.

32. Radulovic E, Mehinagic K, Wüthrich T, Hilty M, Posthaus H et al. (2022) The baseline immunological and hygienic status of pigs impact disease severity of African swine fever. PLoS pathogens 18 (8): e1010522.

33. Baron J, Bin-Tarif A, Herbert R, Frost L, Taylor G et al. (2014) Early changes in cytokine expression in peste des petits ruminants disease. Veterinary research 45 (1): 22.

34. Wani SA, Sahu AR, Khan RIN, Praharaj MR, Saxena S et al. (2021) Proteome Modulation in Peripheral Blood Mononuclear Cells of Peste des Petits Ruminants Vaccinated Goats and Sheep. Frontiers in veterinary science 8: 670968.

35. Wani SA, Praharaj MR, Sahu AR, Khan RIN, Saxena S et al. (2021) Systems Biology behind Immunoprotection of Both Sheep and Goats after Sungri/96 PPRV Vaccination. mSystems 6 (2).

36. Li S (2019) Regulation of Ribosomal Proteins on Viral Infection. Cells 8 (5).

37. Miller CM, Selvam S, Fuchs G (2021) Fatal attraction: The roles of ribosomal proteins in the viral life cycle. Wiley interdisciplinary reviews. RNA 12 (2): e1613.

38. Meng X, Wang X, Zhu X, Zhang R, Zhang Z et al. (2023) Quantitative analysis of acetylation in peste des petits ruminants virus-infected Vero cells. Virology journal 20 (1): 227.

39. 39. Tang J, Tang A, Du H, Jia N, Zhu J, et al. (2022) Peste des Petits Ruminants Virus Exhibits Cell-Dependent Interferon Active Response. Frontiers in Cellular and Infection Microbiology 12: 874936.

40. Li P, Zhu Z, Cao W, Yang F, Ma X et al. (2021) Dysregulation of the RIG-I-like Receptor Pathway Signaling by Peste des Petits Ruminants Virus Phosphoprotein. Journal of immunology (Baltimore, Md. : 1950) 206 (3): 566–579.

41. Miao Q, Qi R, Meng C, Zhu J, Tang A et al. (2021) Caprine MAVS Is a RIG-I Interacting Type I Interferon Inducer Downregulated by Peste des Petits Ruminants Virus Infection. Viruses 13 (3).

42. Chinnakannan SK, Nanda SK, Baron MD (2013) Morbillivirus v proteins exhibit multiple mechanisms to block type 1 and type 2 interferon signalling pathways. PloS one 8 (2): e57063.

43. Rojas JM, Pascual E, Wattegedera SR, Avia M, Santiago C et al. (2021) Hemagglutinin protein of Peste des Petits Ruminants virus (PPRV) activates the innate immune response via Toll-like receptor 2 signaling. Virulence 12 (1): 690–703.

44. Libeau G, Saliki JT, Diallo A (1997) Caractérisation d’anticorps monoclonaux dirigés contre les virus de la peste bovine et de la peste des petits ruminants : identification d’épitopes conservés ou de spécificité stricte sur la nucléoprotéine. Revue Élev. Méd. vét. Pays trop.: 181–190. Available: https://pdfs.semanticscholar.org/2b16/8e0f8701cf39ed93378e86f5bd106a8ce369.pdf.

45. Dobin A, Davis CA, Schlesinger F, Drenkow J, Zaleski C et al. (2013) STAR: ultrafast universal RNA-seq aligner. Bioinformatics (Oxford, England) 29 (1): 15–21.

46. Liao Y, Smyth GK, Shi W (2014) featureCounts: an efficient general purpose program for assigning sequence reads to genomic features. Bioinformatics (Oxford, England) 30 (7): 923–930.

47. Love MI, Huber W, Anders S (2014) Moderated estimation of fold change and dispersion for RNA-seq data with DESeq2. Genome biology 15 (12): 550.

48. Subramanian A, Tamayo P, Mootha VK, Mukherjee S, Ebert BL et al. (2005) Gene set enrichment analysis: a knowledge-based approach for interpreting genome-wide expression profiles. Proceedings of the National Academy of Sciences of the United States of America 102 (43): 15545–15550.

49. Subramanian A, Kuehn H, Gould J, Tamayo P, Mesirov JP (2007) GSEA-P: a desktop application for Gene Set Enrichment Analysis. Bioinformatics (Oxford, England) 23 (23): 3251–3253.

